# Sequence and supercoiling-dependent effects on the structural dynamics of DNA minicircles

**DOI:** 10.1101/2025.08.20.671094

**Authors:** Manuel Micheloni, Luca Tubiana, Raffaello Potestio, Lorenzo Petrolli

## Abstract

The degree of over-/under-winding of the DNA double helix quantified by the superhelical density, is a key feature modulating critical biological processes such as gene expression and regulation: In fact, DNA molecules are able to channel the excess levels of mechanical stress into local defective and denatured states that are promptly detected by, e.g., transcription factors and nuclease enzymes. The occurrence and stability of these motifs is dictated by a complex interplay between topological and sequence-dependent effects, ultimately affecting the global conformational dynamics of the DNA molecule itself. Here, we characterize the impact of the sequence and of the super-helical density on the structural evolution of a 672-bp DNA minicircle *via* classical molecular dynamics simulations employing the coarse-grained oxDNA force field. We observe that moderately-to-highly undercoiled regimes are associated with the occurrence of stable, somewhat broad denaturation bubbles, typically co-localizing with flexible nucleotide sequences on the DNA minicircle: These defects are hardly re-adsorbed by the system, thereby pinning the subsequent dynamics of the molecule. In fact, a similar behavior was recapitulated by enforcing “synthetic”, adjoining DNA mismatches, regardless of the underlying nucleotide sequence, suggesting an effective manner of DNA manipulation.

## Introduction

DNA holds the hereditary information for cells to replicate and develop, and adopts a variety of structural and functional frameworks that vary across species: Key to this polymorphism is the degree of over/underwinding of the DNA double helix about its axis, or supercoiling, which critically affects its structural compartmentalization as well as key cellular processes [1–3]. In prokaryotes, negative supercoiling drives the compaction [4, 5] and local denaturation of DNA, the latter providing a mechanism to modulate the expression of genes [2, 6, 7]. Similarly, eukaryotic DNA exhibits supercoiling emerging from its packing about histone cores in nucleosomes, with topoisomerases regulating the degree of torsional stress through key processes such as DNA transcription and replication [8].

The levels of DNA supercoiling are expressed *via* the superhelical density (*σ*):

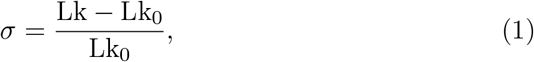

where the linking number (Lk) denotes the number of crossings between the (oriented) strands of a closed, double-stranded DNA molecule, with Lk_0_ defining its reference value (i.e., associated with a torsionally-relaxed conformation) which typically depends upon the thermodynamic environment [9]. The Călugăreanu-White-Fuller theorem expresses the linking number as sum of two contributions, namely Lk = Tw + Wr, with the twist (Tw) quantifying the wrapping of the two strands of DNA about the axis of the double helix, and the writhe (Wr) the coiling of the helical axis in space [10, 11]. In closed dsDNA moieties, the linking number is a topological invariant of the conformational ensemble explored by the system, and is conserved as such [3].

Several efforts have been devoted to characterizing the spontaneous occurrence of DNA defects induced by excess supercoiling levels, as well as the mechanical and biological implications thereof: Coarse-grained, molecular dynamics (MD) simulations of linear DNA molecules in a magnetic-tweezers setup have shown that force-mediated denaturation bubbles typically spawn at the tips of plectonemes (i.e., the apical loop of a coiled DNA structure), and consistently so at AT-rich stretches, acting as channels of stress relief [12]. DNA defects have been observed in regimes of zero external force alike, both by atomistic MD simulations [13] and by experimental assessments of circular DNA molecules [14–16]. Yet, the dynamical interplay between topological and sequence-dependent features—which drive the nucleation of local DNA defects and contributes to reshaping the conformational ensemble of the DNA molecule—has not been explored extensively.

By means of classical MD simulations employing the oxDNA2 coarse-grained force field, here we characterize the impact of the sequence and of excess supercoiling levels on the conformational dynamics of a 672-bp DNA minicircle, as well as their influence on the likelihood and lifetime of denaturation phenomena. DNA minicircles are small, circular dsDNA molecules —less than a few hundred base pairs (bps)—, often employed as structural proxies to complex DNA topologies [1, 17]: On account of their reduced size, the thermodynamical and mechanical behavior of DNA minicircles might be engineered by designing suitable sequences and adjusting the levels of superhelical density [15, 16]—in fact, these systems have been broadly characterized, both experimentally [14–16, 18, 19] and in silico [7, 20–24].

We observe that the amount of (negative) superhelical density majorly affects the likelihood of occurrence of DNA bubbles, with the nucleotide sequence modulating their stability and distribution. In line with earlier works, these defects are most likely associated with AT-rich stretches, on account of their inherent flexibility, and consistently co-localize with apical loops on the DNA molecule [12, 25]: Conversely, DNA bubbles at CG-rich stretches are short (typically involving the failure of a few base pairings) and promptly reabsorbed by the system.

Furthermore, we verify that, while spawning stochastically and homogeneously along the minicircle, denaturation bubbles nucleating at AT-rich stretches often exhibit extended lifetimes—specifically so under highly-negative supercoiling regimes—and exert an effective bias on the molecular structure, thereby pinning the conformational dynamics of the minicircle (in line with earlier experimental data [15]). This behavior is recapitulated by placing multiple adjoining DNA mismatches along the sequence: In fact, DNA defects as small as 5 bps are capable to disrupt the equilibrium dynamics of the minicircle by forcing the migration of (otherwise stable) supercoiled regions into an apical loop, suggesting an effective manner of structural and topological DNA manipulation, e.g., enhancing such processes as DNA binding and recognition.

## Materials and Methods

### System setup and simulation protocol

The 672-bp circular DNA molecule employed in this work matches the template reported by Fogg and co-workers [15], who joined the sequence of a known 336-bp DNA minicircle in a tandem orientation (see Section 1 of the Supplementary Material for the full DNA sequence and Figure 1 showing the density of AT/CG pairings along the DNA template).

**Figure 1:**
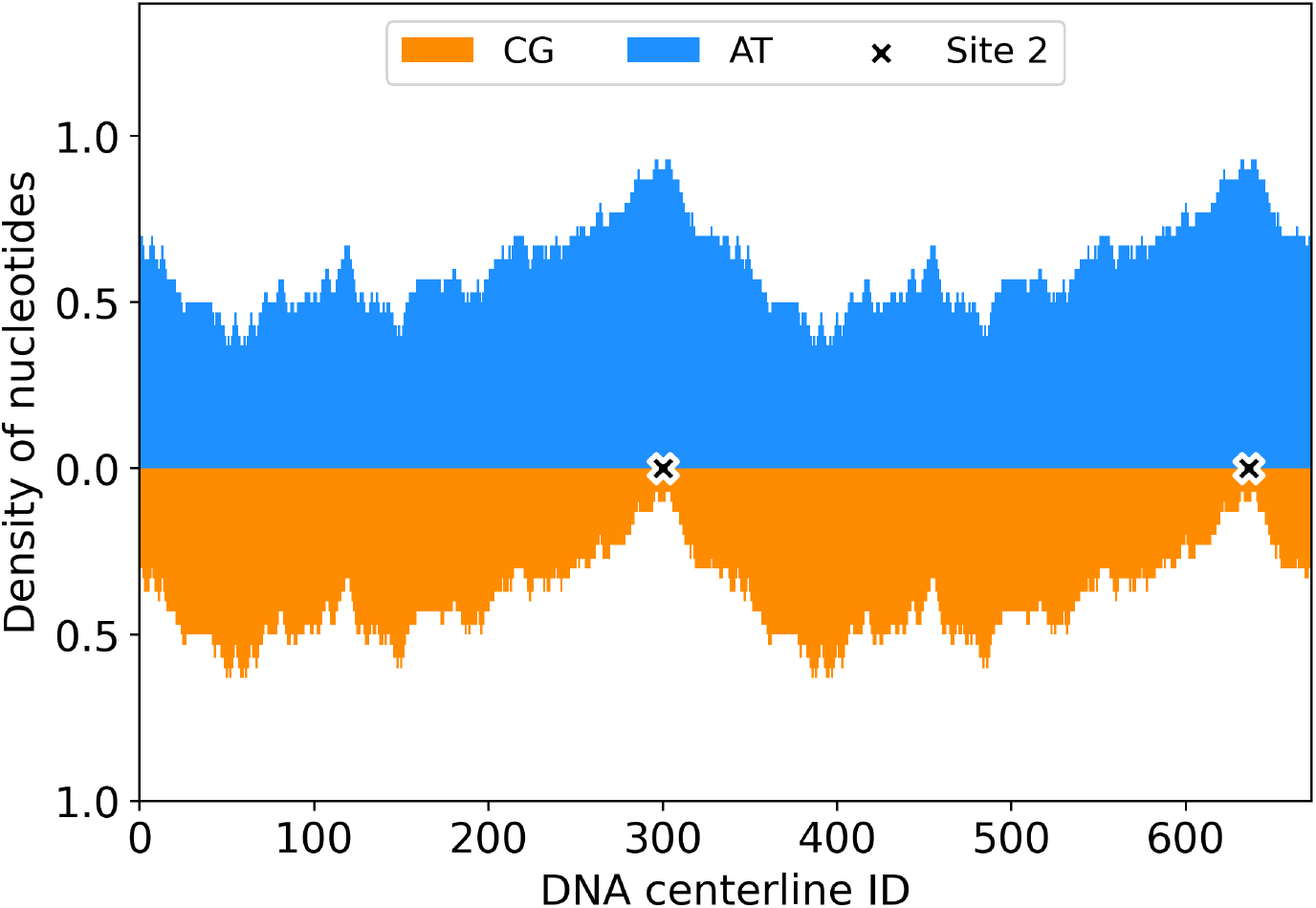
(Rolling average of) The AT/CG density along the centerline of the 672-bps DNA minicircle employed in this work, based upon the template sequence of Fogg and co-workers [15] (reported in Section 1 of the Supplementary Material). The location of the cleavage sites of the Bal-31 nuclease according to Ref. [15] (site 2) is identified by small icons.

Six degrees of superhelical density have been chosen—namely *σ* = −0.1, −0.08, −0.06, 0, 0.06 and 0.1—to assess the impact of the topological regime on the conformational dynamics of the DNA minicircle. All simulations have been performed employing the oxDNA2 CG force field [26] taking into account sequence-specific hydrogen-bonding and stacking interactions. Ten independent MD replicates have been carried out for each level of superhelical density, at a physiological monovalent salt concentration of 0.15 M (which oxDNA2 models effectively *via* a Debye-Hückel electrostatic screening) and temperature T = 310 K, employing the GPU-accelerated, native oxDNA code [27, 28]. Particularly, a three-step MD protocol has been followed in each replicate:

- The (perfectly circular) starting configuration of the DNA minicircle is generated *via* tacoxDNA [29]: For each supercoiling regime, a proper topological constraint is enforced by fixing the corresponding value of *σ*, and homogeneously distributing the twist imbalance along the DNA filament;
- Subsequently, the DNA minicircle is subjected to a swift relaxation stage of 1 *×* 10^7^ MD steps *via* the Bussi thermostat [30], associated with a timestep Δ*t* = 9.09 fs (details of the conversion factor between oxDNA and SI units are reported in Ref. [31]). As suggested by the oxDNA developers, the correlation time and the timestep frequency of the thermostat have been set to 1000 and 53 MD steps respectively. Moreover, at this stage, a modified version of the backbone potential [32] has been enabled, ensuring a higher stability towards the larger displacements of the CG sites;
- A production run of 5 *×* 10^8^ MD steps (associated with a sampling frequency of 5 *×* 10^5^ MD steps) is thus performed starting from the relaxed DNA conformation, employing the Langevin thermostat: The diffusion coefficient and timestep frequency of the thermostat have been set to 2.5 and 103 MD steps respectively, according to the recommendations of the oxDNA developers [31]. The overall sampling time per replicate has been established upon (the decay of) the autocorrelation of the gyration radius of the DNA minicircle at the diverse degrees of super-helical density—refer to Section 2 of the Supplementary Material for details.

### Structural analysis of the DNA minicircle

Circular DNAs attain diverse configurations upon the seamless interconversion of twist and writhe, driven by the degree of superhelical density [33]. In fact, the dynamical equilibrium of twisted and writhed conformations is fairly well reproduced by the oxDNA2 force field, whereby the DNA mini-circles feature sharply-bent apical loops and highly-intertwined, supercoiled tracts (see Figure 2).

**Figure 2:**
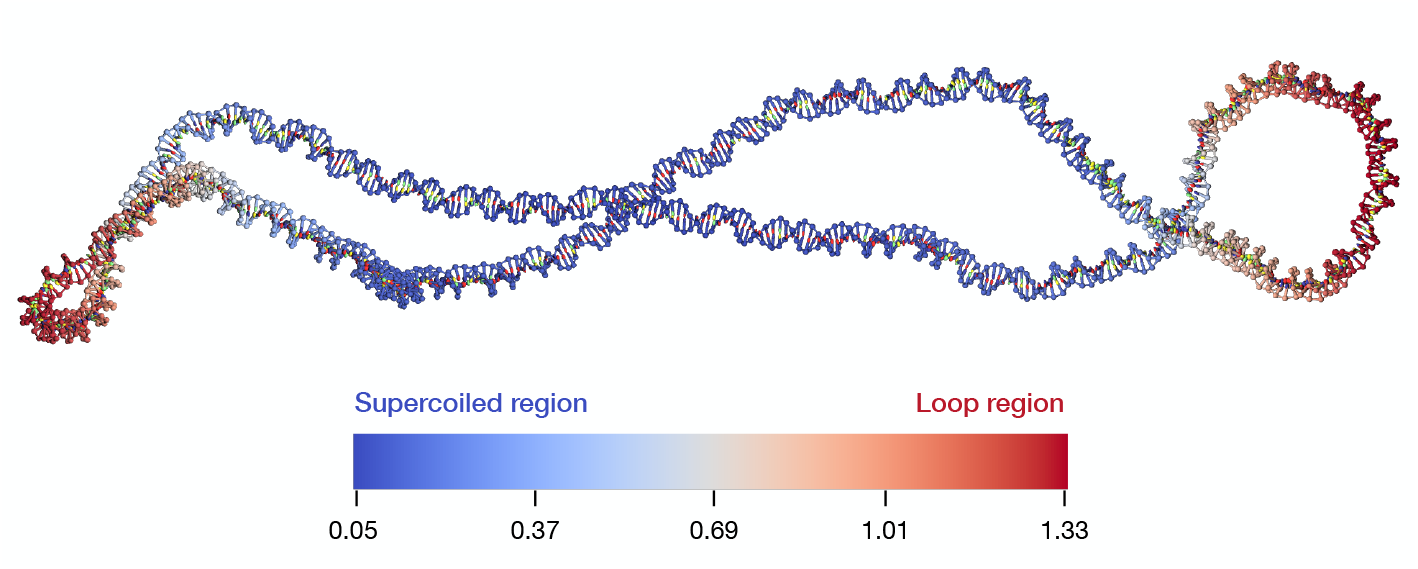
Relaxed conformation of a DNA minicircle at superhelical density *σ* = −0.06, the color bar defined upon the value of the local writhe about the *i*-th nucleotide. Two regions of topological significance are highlighted, i.e., the apical loops and a broad supercoiled tract, the latter characterized by a highly-intertwined DNA bundle.

The evolution of loops and supercoiled tracts along the MD replicates is tracked *via* the (unsigned) *local writhe* |*ω*(*i, t*) | [34, 35] (or the average crossing number, ACN [36]), defined as:

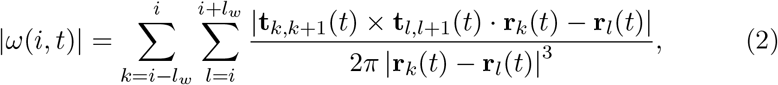

with **r**_*i*_ the coordinate vector of the *i*-th nucleotide pairing along the center-line of the DNA helix, **t**_*i*,*i*+1_ the tangent vector to the helical axis at location *i*, and *l*_*w*_ = 150 bps—corresponding to the persistence length of DNA in solution. From this definition, it follows that a straight DNA segment centered at location *i* contributes zero to the summation in Equation 2, whereas local maxima of |*ω*(*i, t*) | along the helix centerline are associated with highly-bent DNA tracts and/or apical loops. An example calculation of the local writhe on a DNA minicircle at *σ* = −0.06 (and the topologically-significant regions thereof) is shown in Figure 2, while Figure S3 of the Supplementary Material depicts the time evolution of |*ω*(*i, t*)| along a MD replicate at *σ* = −0.06.

## Results and Discussion

### Supercoiling- and sequence-dependent effects on the dynamics of DNA minicircles

We approached our characterization of the conformational dynamics of DNA minicircles by tracking the distribution and stability of the apical loops at the diverse degrees of superhelical density. Figure 3A shows the time average (over all independent MD replicates) of the local writhe (⟨|*ω*(*i*)|⟩) along the DNA centerline: Higher values of |*σ*| are associated with higher values of ⟨|*ω*(*i*)|⟩, as well as with a higher propensity of apical loops to co-localize with specific segments of the DNA sequence. In fact, a few observations are in order:

**Figure 3:**
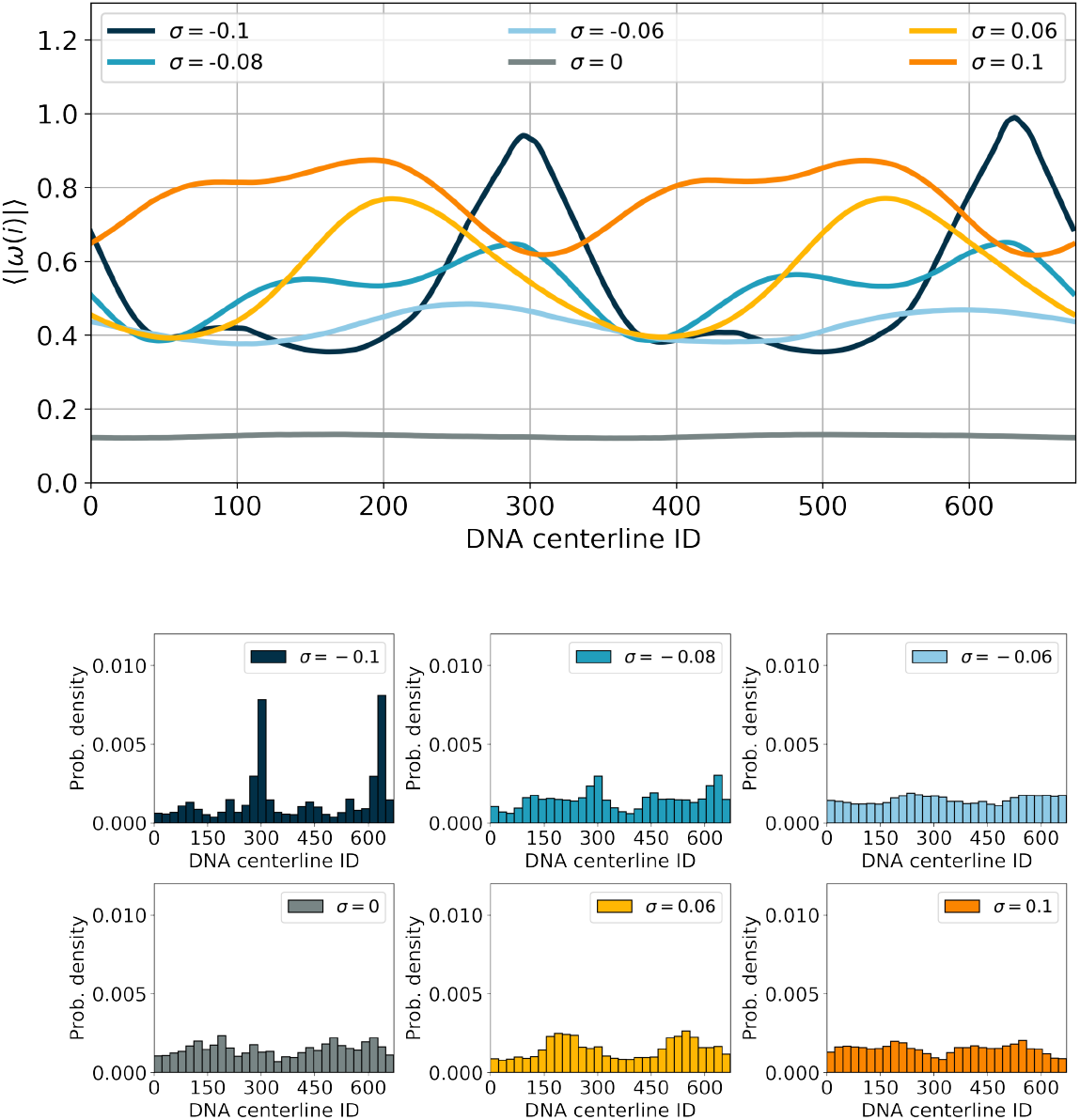
(**Top**) Time average (over all independent MD replicates) of the local writhe (⟨|*ω*(*i, t*)|⟩ along the DNA centerline, for each superhelical density regime. (**Bottom**) Histograms depicting the probability distribution for an apical loop to localize about a specific nucleotide (index) on the DNA minicircle, at the diverse levels of superhelical density.

- at all degrees of *σ*, the values of the average (unsigned) local writhe is replicated symmetrically between the conjoined 336-bps DNA sequences in the 672-bps template, highlighting a robust substrate selectivity in the distribution of apical loops;
- the maxima of ⟨|*ω*(*i*)|⟩ are somewhat consistent between scenarios belonging to either positively- (*σ >* 0) or negatively-supercoiled (*σ <* 0) regimes, and co-localize with the AT-richest DNA segments in the latter case (see Figure 1);
- unlike any other scenario, the values of ⟨|*ω*(*i*)|⟩ at *σ* = − 0.1—as well as the probability distribution of apical loops along the DNA centerline (shown in Figure 3B)—are sharply peaked about specific locations on the minicircle, on account of local DNA denaturation phenomena (*vide infra*). Notably, this behavior is in line with the experimental observations of Fogg and co-workers [15], who mapped the selective cleavage of DNA minicircles at *σ* = −0.096 from the digestion patterns of the Bal-31 nuclease and onto the nucleotides corresponding to a centerline index of about 300 and 600 (as shown in Figure 1).

### Supercoiling- and sequence-dependent effects on DNA denaturation bubbles

As briefly mentioned earlier, the behavior shown by ⟨|*ω*(*i*)|⟩ at highly-negative regimes of superhelical density is accounted for by the occurrence of local denaturation bubbles—i.e., the metastable dissociation of the DNA helix driven by the local failure of the hydrogen bonding interactions (see Figure 4). We hereby define an effective free energy difference Δ*G*_BF_, associated with the likelihood that each topoisomer spawns a denaturation bubble, as:

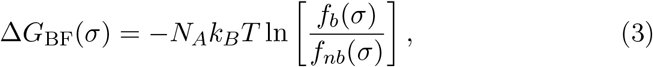

with *k*_*B*_ the Boltzmann constant, *N*_*A*_ the Avogadro’s number and *f*_*b*_, *f*_*nb*_ the fraction of frames in the dynamics of a DNA minicircle exhibiting bubbles or the absence thereof, respectively.

**Figure 4:**
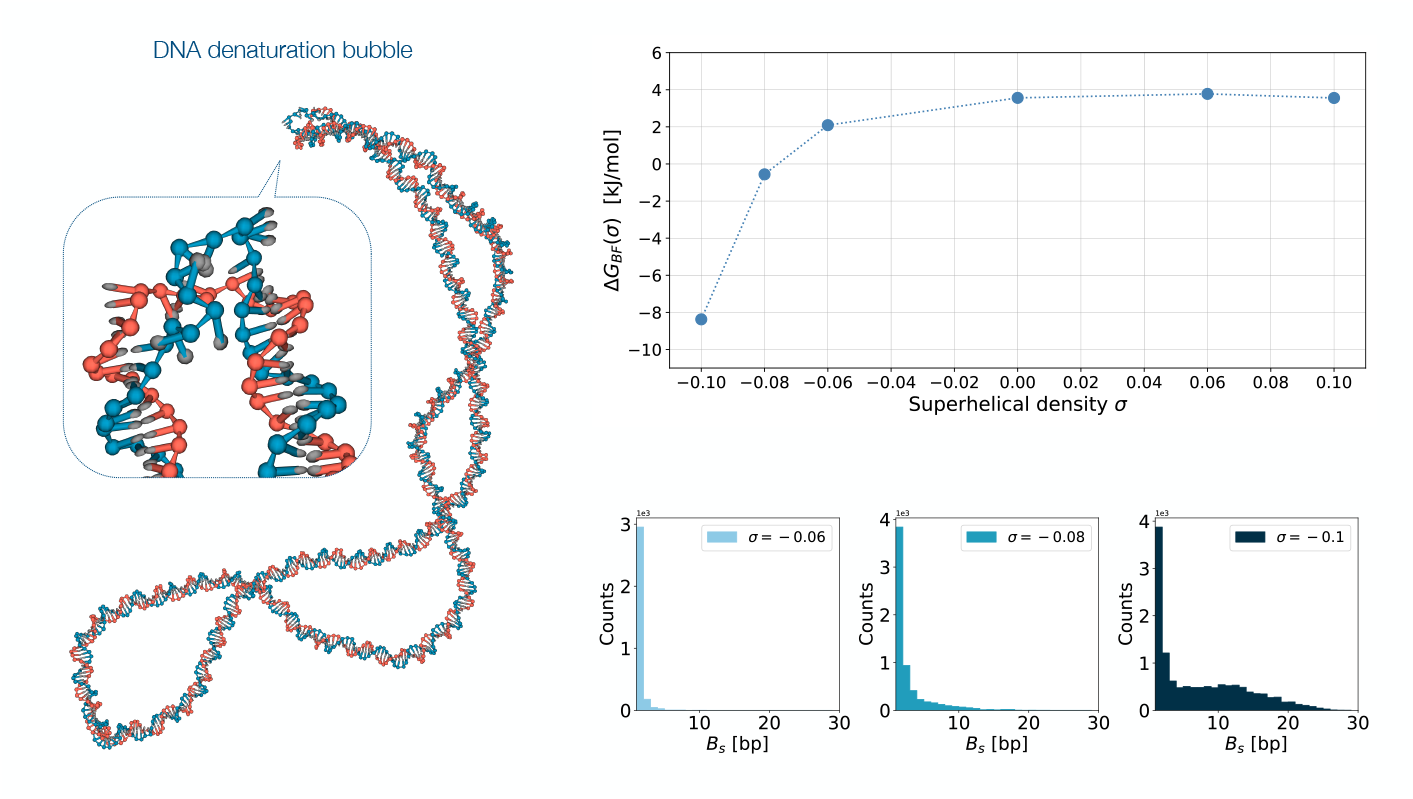
(**Left**) MD snapshot of a DNA minicircle at superhelical density *σ* = −0.1 exhibiting a DNA denaturation bubble (inset). (**Top, right**) Profile of the effective free energy Δ*G*_BF_(*σ*), associated with the propensity of the DNA minicircle to spawn denaturation bubbles as function of the superhelical density—details in the main text and in the Supplementary Material. (**Bottom, right**) Distribution of bubble sizes under negative supercoiling regimes, associated with values of ⟨*B*_*s*_⟩ = 1.1 *±* 0.6 (*σ* = −0.06), 2 *±* 2 (*σ* = −0.08), and 7 *±* 6 (*σ* = −0.1) failed base pairings respectively.

The likelihood of occurrence of DNA bubbles as function of the superhelical density, and the free energy profile Δ*G*_BF_, are shown in Table S1 of the Supplementary Material and in Figure 4 illustrating the response of the minicircles subjected to the diverse supercoiling regimes.

The inherent propensity of negatively supercoiled DNA molecules to yield denaturation bubbles is consistent with the observation that these topological regimes facilitate the strand separation, which is know to enhance sequence recognition and DNA binding processes [37–39]. Particularly, values of *σ* between −0.08 and −0.06 seemingly mark a transition between two thermodynamic regimes, i.e., either favoring (*σ <* −0.07 [15]) or inhibiting the spontaneous occurrence of DNA bubbles. The behavior shown in Figure 4 agrees with the experimental data of Fogg and co-workers, who observed that the topoisomers of the 672-bp minicircle exhibiting highly-negative supercoiling levels are cleaved promptly by Bal-31 [15]: In fact, the frequency and size distribution of the DNA denaturation phenomena recapitulate the cleavage selectivity of the nuclease enzymes, with Bal-31 and S1 binding shorter (single-nucleotide) and larger (≥ 4-nucleotide) bubbles respectively [15].

Conversely, under both zero and positive supercoiling regimes, the likelihood of denaturation events is lower and mostly ascribable to single-nucleotide phenomena: This is in line with the experimental data of Irobalieva and co-workers, based on the enzymatic activity of Bal-31 on 336-bps minicircles at ΔLk = +3 (*σ* = 0.093) [16].

As similarly inferred from the distribution and stability of looped and supercoiled regions, denaturation bubbles favorably co-localize with flexible sequences along the DNA centerline, and consistently so in moderately-to-highly undercoiled regimes: In fact, the occurrence of bubbles is broadly favored at AT-rich DNA stretches, as shown in earlier works alike [19, 40, 41], on account of their inherent capability to bear excess torsional stresses. A significant correlation exists between the (average) locations of apical loops and denaturation bubbles—as shown by Figure 5 and Figure S4 of the Supplementary Material—in line with earlier AFM measurements [14, 42], wet-lab experiments [15, 16, 43], and numerical simulations [12, 40]. Consistently, the distribution of denaturation phenomena peaks at diametrically-opposed sites of the DNA centerline, i.e., about nucleotides 300 and 600 (i.e., the tandem cleavage sites shown in Figure 1), which Fogg and co-workers identified as the sole substrates of the action of the Bal-31 nuclease [15]. Of note, the off-diagonal datapoints shown in Figure 5 and observed at highly-negative supercoiling regimes (*σ* ≤ − 0.08) are associated with DNA bubbles spontaneously spawning about nucleotides ∼300 and ∼600, but not perfectly co-localizing with the apical loops closeby. In fact, we verified that, if these bubbles are sufficiently stable, they drive the subsequent translocation of the loops along the DNA centerline (and about nucleotides ∼300 and ∼600), ultimately “pinning” the dynamics of the DNA minicircle.

**Figure 5:**
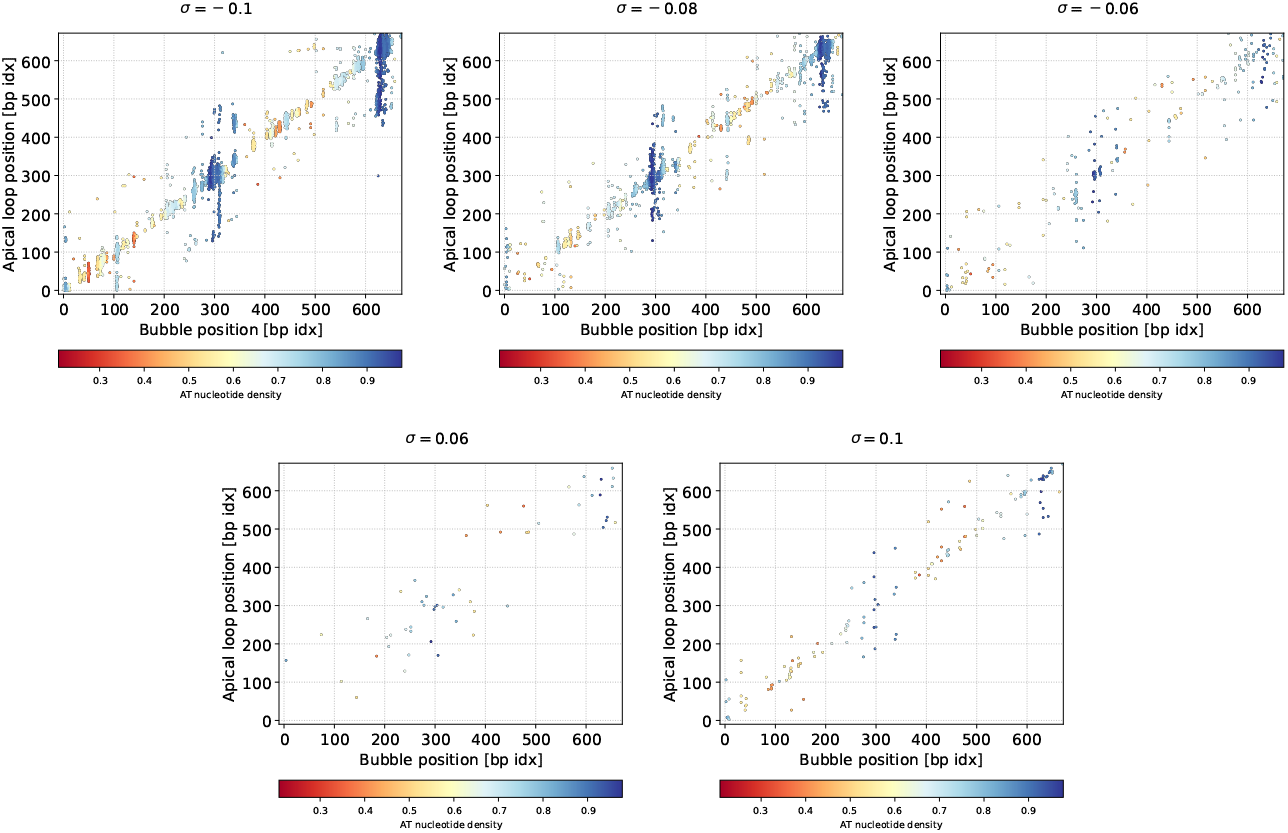
Scatter plot highlighting the correlation between the location of apical loops and of DNA bubbles over all independent MD replicates, at varying degrees of superhelical density: The color code is associated with the local AT density upon a rolling average of 10 nucleotides. For the purpose of clarity, denaturation bubbles of size *B*_*s*_ = 1 are not shown—in fact, they are homogeneously distributed along the DNA centerline.

### Lifetime of DNA denaturation bubbles

We have shown that the likelihood and stability of denaturation bubbles is markedly affected by the sequence and by topological constraints, favoring their co-localization with AT-rich DNA stretches and apical loops—and consistently so at highly-negative supercoiling regimes. Particularly, this bears an impact on both the size distribution and the lifetime of the bubbles, as shown by Figure 6 and Figure S5 of the Supplementary Material: In fact, their average dwelling times (⟨Δ*τ* (*i*)⟩) vary significantly between the diverse regimes of (negative) supercoiling and along the DNA centerline, with the DNA bubbles co-localized with AT-rich stretches exhibiting the highest life-times.

**Figure 6:**
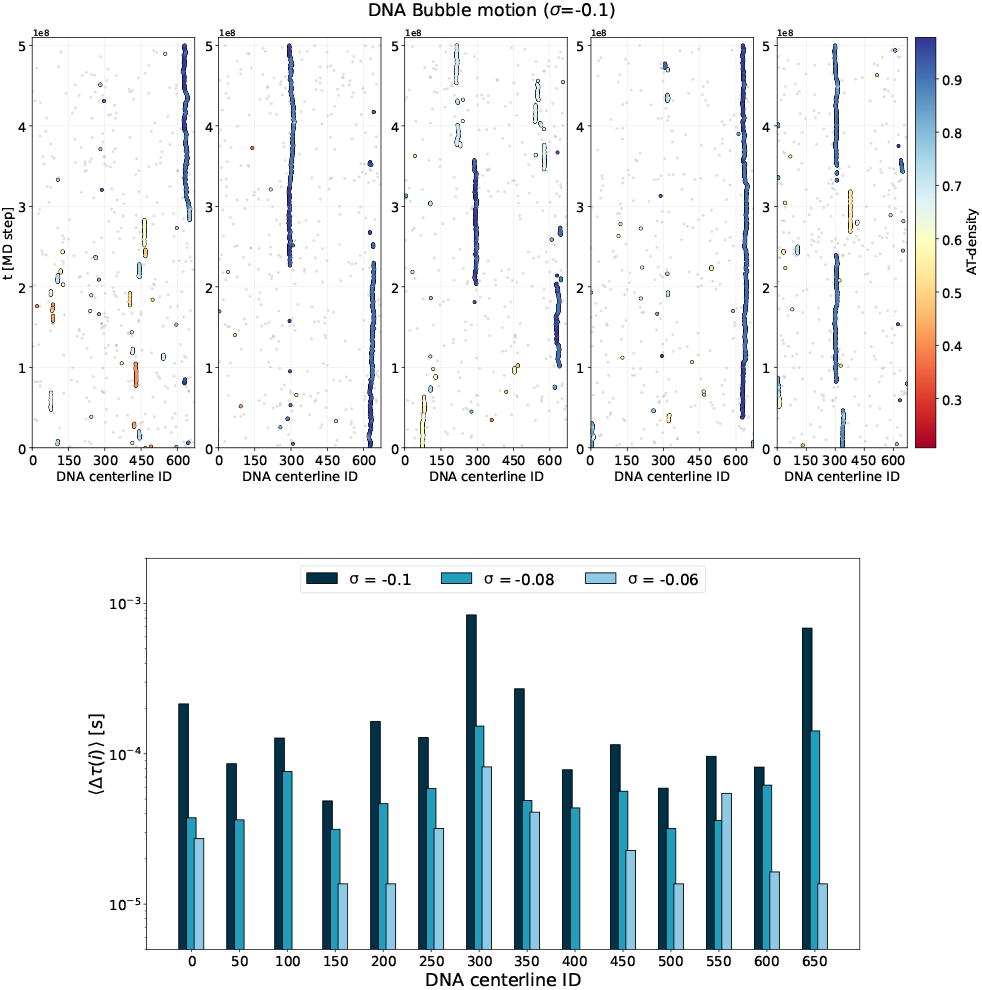
(**Top**) Time evolution of denaturation bubbles along the centerline of a DNA minicircle: Each data-point is defined by the center of mass of the bubble lying (approximately) about the *i* -th nucleotide. A color scale is attributed to each bubble upon the local AT density (over a rolling average of 10 nucleotides) but for the denaturation of single base pairings—which are depicted in gray for the purpose of clarity. Here, the outcome of five independent MD simulations at superhelical density *σ* = −0.1 is shown from left to right. (**Bottom**) The average lifetime of denaturation bubbles about each nucleotide of the DNA minicircle is shown in logarithmic scale, at diverse degrees of (negative) superhelical density. For the purpose of statistical significance, the values of ⟨Δ*τ* (*i*)⟩ have been grouped by windows of 50 nucleotides about a reference index *i*. Likewise, all events associated with the failure of single base pairings have been neglected in the calculation.

It is well acknowledged that the integration of multiple degrees of freedom associated with the coarse-graining procedure smoothens the free-energy landscape of a system (with respect to their atomistic counterpart), thereby enhancing its dynamics and favoring the transitions between thermodynamic basins [44–46]. Moreover, the oxDNA force field is bundled with an implicit description of the solvent, thus lacking a proper characterization of hydrodynamic effects [47, 48]. In light of their kinetic implications, a rescaling of the time steps of the CG simulation is typically performed upon an atomistic MD benchmark or wet-lab experiment, under the approximation that the modifications induced by the model are quantifiable in terms of a global redefinition of the characteristic times of a system [44].

By assuming that a DNA bubble might be described as a local melting/hybridization process, a rescaling factor of 3000 has been employed to convert internal into SI time units [49], based on the agreement between the oxDNA simulations and the experimental rates of the DNA hybridization [26].

Notably, our lifetime estimates are consistent with the values reported by Altan-Bonnet and co-workers, who inferred the characteristic timescales of bubble (un)zipping ranging between 20 - 100 *µs* [50] (see Figure 6). Moreover, we observe that the shape of the bubbles seamlessly adjusts to the DNA sequence (i.e., with the bubble sliding across the centerline), e.g. by broadening about AT-rich DNA stretches—as shown by Figure 7 and Figure S6 of the Supplementary Material: This tallies with the work of Hwa and co-workers, who characterized the dynamics of twist-induced bubbles in random DNA sequences, showing that smaller defects are distributed ubiquitously along the DNA sequence, whereas larger bubbles co-localize consistently with AT-rich DNA stretches [51]. Additionally, higher (negative) supercoiling regimes favor the occurrence of larger defects that are hardly reabsorbed by the system, which accounts for their extended lifetime.

**Figure 7:**
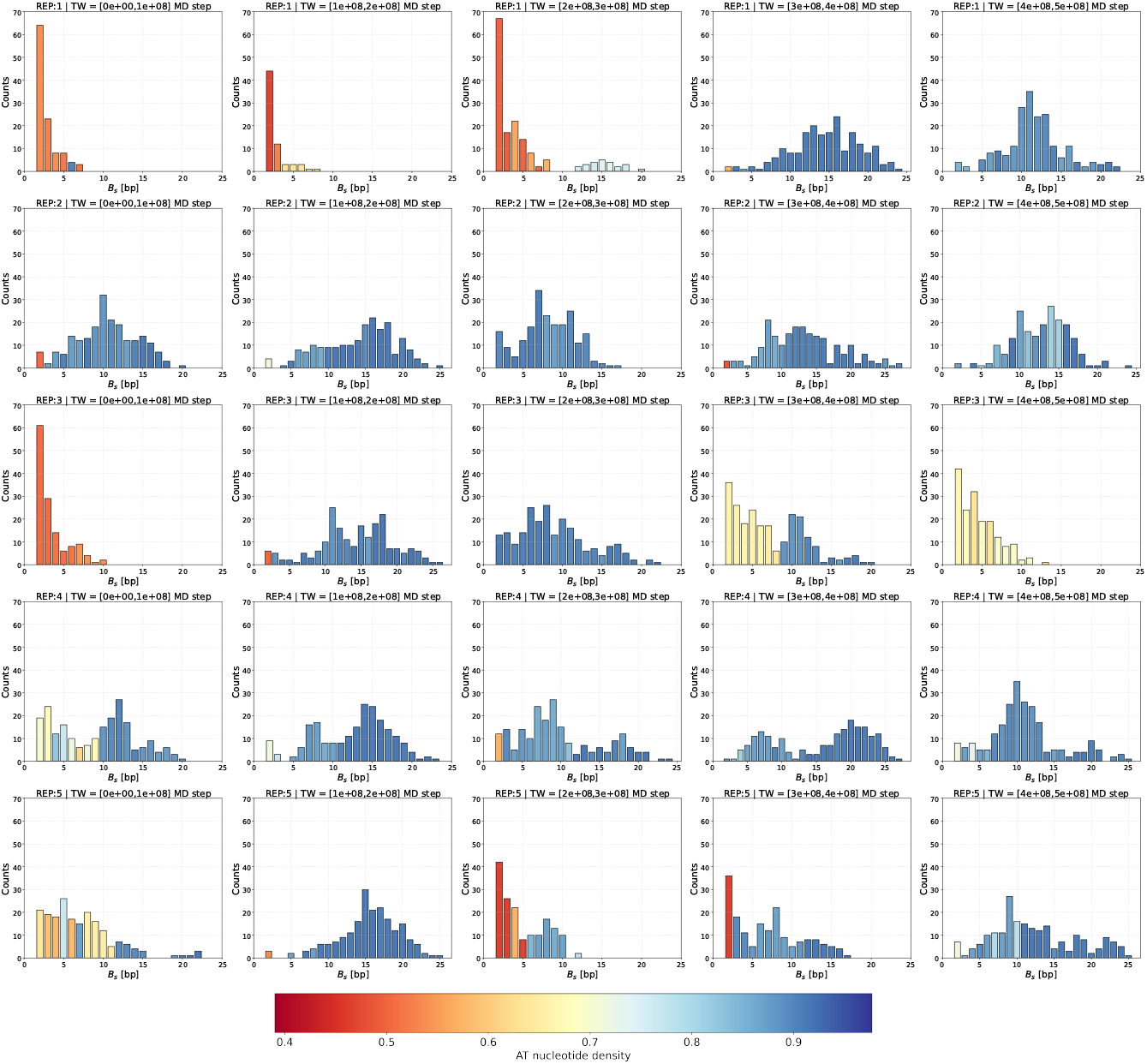
Histograms showing the distribution of DNA bubble sizes within time windows (TW) of 1*×*10^8^ MD steps, along a selection of MD replicates in the *σ* = −0.1 scenario (corresponding to the trajectories shown in Figure 6. A color code has been defined upon (the statistical mode of) the distribution of AT-density values co-localizing with bubbles of a specific size. For the purpose of clarity, all events involving the failure of single nucleotide pairings have been neglected.

### Bubbles drive the structural dynamics of DNA minicircles

We have described the process whereby denaturation bubbles affect the dynamics of DNA minicircles by migrating and co-localizing with apical loops. In fact, this was similarly observed *in silico* by Desai and co-workers, who inferred that DNA mismatches (MMs) beyond a specific size pin the dynamics of a DNA plectoneme—within a magnetic tweezer setup, and under specific regimes of supercoiling, external force and saline concentration [52].

Here, we further characterize this phenomenon at a negative degree of superhelical density, by enforcing (adjoining) mismatches upon a DNA minicircle at *σ* = −0.06 as “synthetic bubbles” of size *B*_*s*_ = 1, 2, 5, 10 bps: To this purpose, we have selectively replaced nucleotide-pairings along the DNA sequence with special nucleotides [53] that retain all interactions of the oxDNA force field but the Watson-Crick hydrogen bonding (see Figure 8 and refer to the oxDNA documentation for details). We shall remark that, although mismatched nucleotides establish (weak) non-canonical interactions [54], we will hereby assume that their significance is negligible in this context. Moreover, to try and avoid any interference with spontaneous denaturation events, all synthetic bubbles have been placed on supercoiled regions of the minicircle showing a low denaturation propensity in the equilibrated trajectories (see Table S2) of the Supplementary Material).

**Figure 8:**
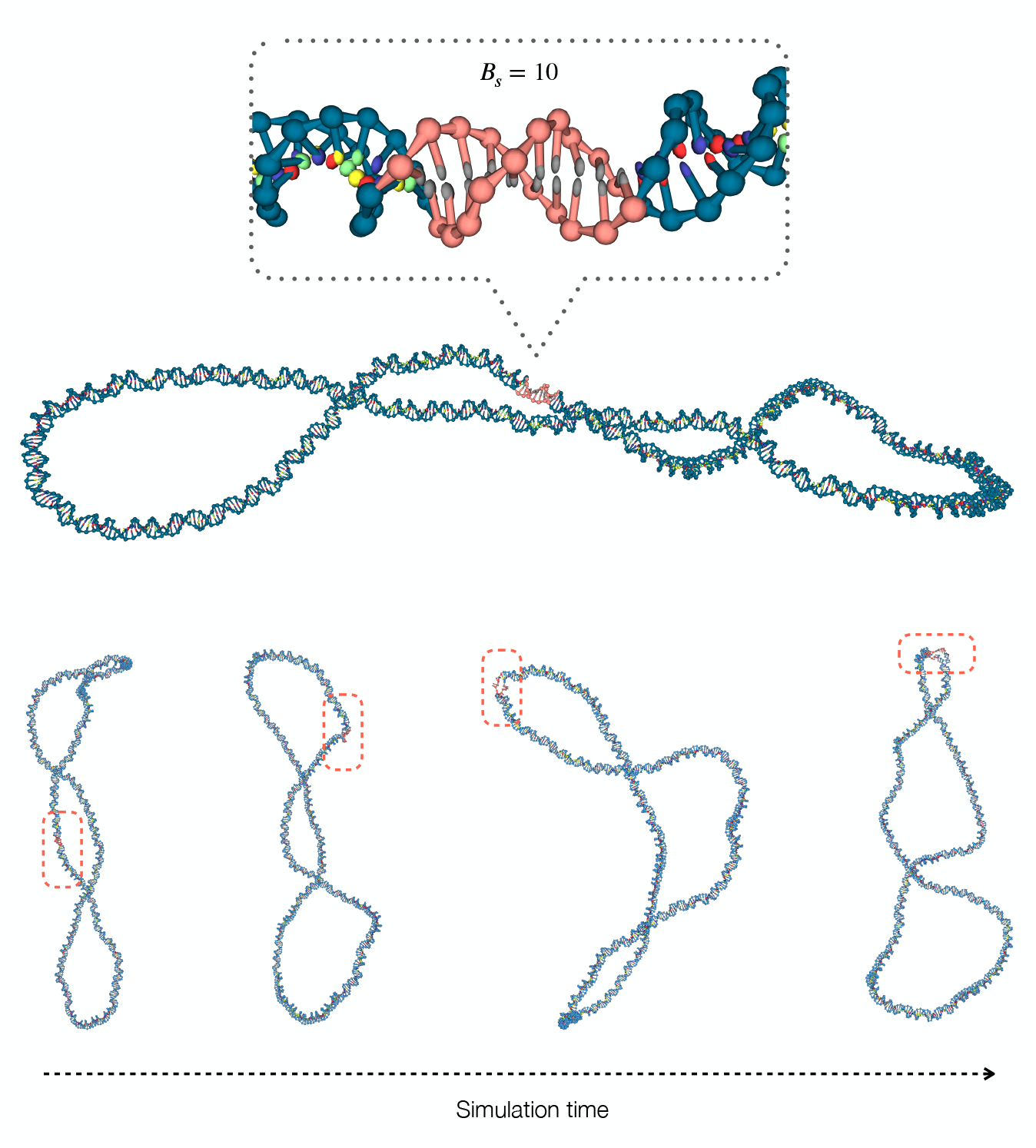
(**Top**) A synthetic bubble of size *B*_*s*_ = 10 bps is enforced on the supercoiled region of a 672-bp DNA minicircle characterized by a superhelical density *σ* = −0.06. (**Bottom**) The concerted, structural adjustment of a DNA minicircle driven by a synthetic bubble of size *B*_*s*_ = 5 placed along the supercoiled region, whereby the defect steadily migrates towards an apical loop.

Overall, the impact of synthetic bubbles is proportional to the extent of the mismatch, and affects the behavior of the DNA minicircle on both a structural and dynamical level: In fact, Figure 9 shows that the propensity of DNA mismatches to migrate and co-localize with apical loops increases with the size of the defect (an example process is shown in Figure 8).

**Figure 9:**
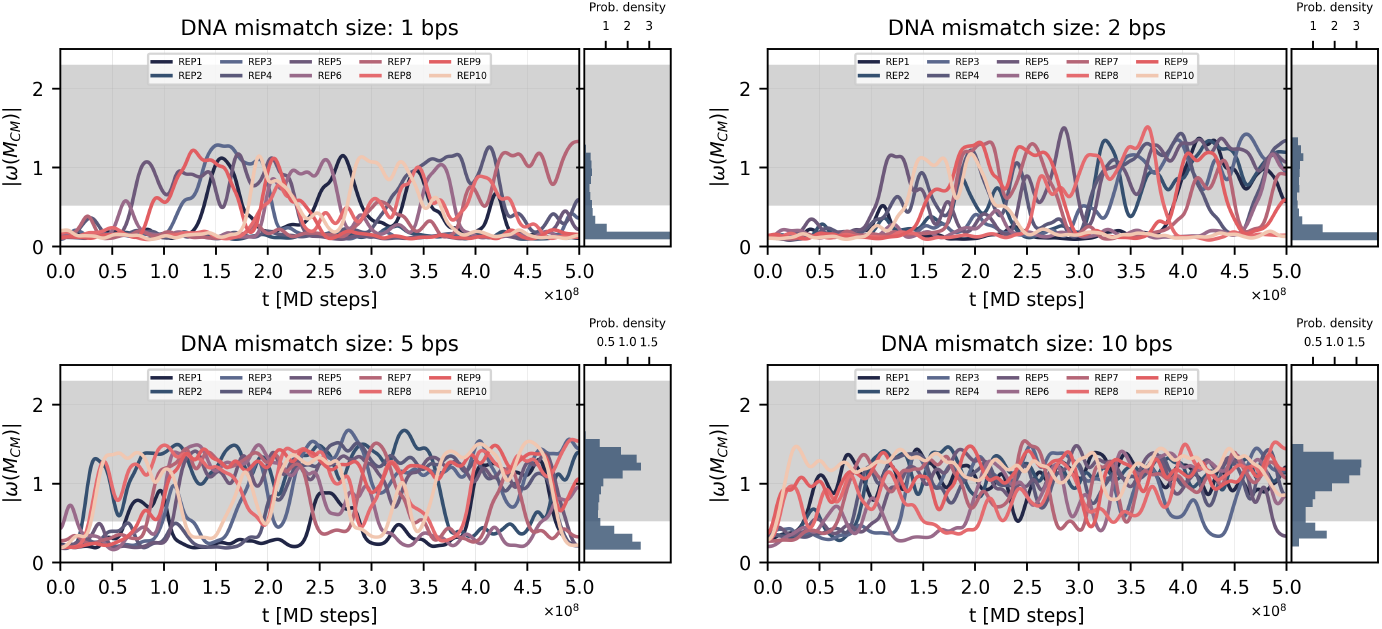
Time evolution of the (unsigned) local writhe about the centroid of a synthetic bubble *B*_*CM*_ (and the distribution thereof), upon diverse MD replicates and bubble sizes *B*_*s*_ = 1, 2, 5, 10 bps. The shaded area highlights the distribution of |*ω*(*i, t*)| values associated with the apical loop regions of a DNA minicircle at equilibrium and superhelical density *σ* = −0.06 (refer to Figure S7 of the Supplementary Materials).

From a quantitative standpoint, we express this propensity in terms of the ratio *κ* between the average dwelling time of a synthetic bubble at an apical loop (l) and at the supercoiled region (sc) as:

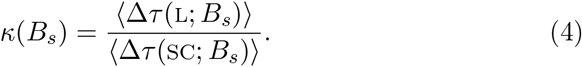

he values of *κ* are summarized in Table 1 and benchmarked against a characteristic *κ*_*eq*_—defined by the propensity shown by each nucleotide of the DNA sequence to dwell about either (l, sc) region of the minicircle at equilibrium and superhelical density *σ* = −0.06. This choice is justified by the absence of a significant pinning effect along the sequence at *σ* = −0.06 (see Figure 3), such that all nucleotides show an equal probability to host an apical loop, on average. In fact, the dwelling times of nucleotides at equilibrium are 4.4 *×*10^6^ and 1.2 *×*10^7^ MD steps in loop and supercoiled regions respectively, with a value of *κ*_*eq*_ = 0.3. As shown in Table 1, the increase of *κ* with *B*_*s*_ highlights the capability of small synthetic bubbles (*B*_*s*_ = 5, 10) to bias the structural dynamics of the minicircle at equilibrium—whereas the impact of point-like defects is less significant on a global scale.

**Table 1:** Average dwelling times of the synthetic bubbles at apical loops (l) and at the supercoiled (sc) region of a DNA minicircle at *σ* = −0.06, and the ratio thereof (*κ*). The reference value *κ*_*eq*_ = 0.3 is estimated upon the equilibrium dynamics of the minicircle at *σ* = −0.06, as detailed in the main text.

| $B_s$ [bp] | $\langle \Delta\tau(\text{SC}) \rangle$ [MD steps] | $\langle \Delta\tau(\text{L}) \rangle$ [MD steps] | $\kappa$ | $\kappa/\kappa_{eq}$ |
| --- | --- | --- | --- | --- |
| 1 | $2.1 \times 10^7$ | $4.8 \times 10^6$ | 0.2 | 0.67 |
| 2 | $1.6 \times 10^7$ | $5.6 \times 10^6$ | 0.3 | 1 |
| 5 | $7.0 \times 10^6$ | $1.2 \times 10^7$ | 1.7 | 5.67 |
| 10 | $3.0 \times 10^6$ | $1.0 \times 10^7$ | 3.4 | 11.33 |

## Conclusions

The degree of superhelical density is a critical feature driving the (mechanical, thermodynamical) stability of the DNA double helix, and plays diverse roles in the regulation, expression, and organization of the genome. In this work, we have characterized the structural dynamics of a 672-bp DNA mini-circle subjected to diverse levels of supercoiling, *via* a broad MD simulation campaign with the oxDNA force field. Specific efforts have been devoted to assess the correlations between the sequence, the topological constraints of the DNA molecule (defined by the superhelical density), and the likelihood and stability of local DNA defects, such as denaturation bubbles. In fact, the release of the excess mechanical stress associated with moderate-to-high levels of negative supercoiling occurs through local strand separation phenomena—induced by the failure of hydrogen bonds—which themselves trigger a global conformational adjustment of the DNA molecule.

Particularly, major denaturation events characterized by an extended life-time spontaneously occur at *σ* = −0.08 and −0.1, and consistently co-localize with AT-rich DNA stretches at apical loops—thereby enforcing a bias on the conformational ensemble of the minicircle. To this concern, we notice that, despite the nucleation of bubbles being an inherently stochastic phenomenon, the degree of supercoiling and the DNA sequence ultimately modulate the structural dynamics of the DNA minicircle.

The influence of denaturation defects has been further verified by enforcing non-reversible, “synthetic” bubbles (as single or adjoining DNA mismatches) on the supercoiled region of the DNA minicircles. Notably, beyond a threshold bubble size ≥ 5 base pairings, these defects significantly bias the conformational ensemble of DNA minicircles, forcing the translocation of a target DNA stretch into an apical loop, regardless of the nucleotide sequence and the equilibrium dynamics thereof. In fact, a similar behavior is attributable to spontaneous bubbles alike, despite major denaturation events rarely spawning but at higher levels of DNA supercoiling.

In conclusion, these observations emphasize the role of the dynamical interplay between the DNA sequence and topology, as a critical factor in biological processes such as sequence recognition and DNA binding, and suggest an effective manner of structural DNA manipulation by design.

## Author Contributions

LP and RP oversaw and coordinated the whole research project. MM performed all MD simulations. MM, LP and LT designed the analysis protocol: The data analysis was carried out by MM with the constant supervision of LP. MM and LP mainly worked on the first draft, while the final manuscript was written with contributions from all authors.

## Supplementary Material

Supplementary Material is available. Moreover, the raw data associated with this work are freely available on a Zenodo repository at the link: https://zenodo.org/records/16648999.

## Acknowledgments

The authors acknowledge support from the following Agencies: CINECA award under the ISCRA initiative, for the availability of high-performance computing resources; Fondazione CARITRO through the project COMMODORE (#20260); the Italian Ministry of Education, University and Research (MIUR) through the FARE grant for the project HAMMOCK (grant R18ZHWY3NC); ICSC – Centro Nazionale di Ricerca in HPC, Big Data and Quantum Computing, funded by the European Union under NextGenerationEU; PRIN 2022 PNRR Prot. n. 2022Z3BBPE. Views and opinions expressed are however those of the author(s) only and do not necessarily reflect those of the European Union or The European Research Executive Agency. Neither the European Union nor the granting authority can be held responsible for them.

