## Supplementary Material for "Sequence and supercoiling-dependent effects on the structural dynamics of DNA minicircles"

#### 1 DNA template

Here is reported the 5'-3' sequence of the 672-bps DNA minicircle employed in this work, based on the template of Fogg and co-workers [1].

```
  1 TTTATACTAA CTTGAGCGAA ACGGGAAGGG TTTTCACCGA TATCACCGAA
 51 ACGCGCGAGG CAGCTGTATG GCATGAAAGA GTTCTTCCCG GAAAACGCGG
101 TGGAATATTT CGTTTCCTAC TACGACTACT ATCAGCCGGA AGCCTATGTA
151 CCGAGTTCCG ACACTTTCAT TGAGAAAGAT GCCTCAGCTC TGTTACAGGT
201 CACTAATACC ATCTAAGTAG TTGATTCATA GTGACTGCAT ATGTTGTGTT
251 TTACAGTATT ATGTAGTCTG TTTTTTATGC AAAATCTAAT TTAATATATT
301 GATATTTATA TCATTTTACG TTTCTCGTTC AGCTTTTTTA TACTAACTTG
351 AGCGAAACGG GAAGGGTTTT CACCGATATC ACCGAAACGC GCGAGGCAGC
401 TGTATGGCAT GAAAGAGTTC TTCCCGGAAA ACGCGGTGGA ATATTTCGTT
451 TCCTACTACG ACTACTATCA GCCGGAAGCC TATGTACCGA GTTCCGACAC
501 TTTCATTGAG AAAGATGCCT CAGCTCTGTT ACAGGTCACT AATACCATCT
551 AAGTAGTTGA TTCATAGTGA CTGCATATGT TGTGTTTTAC AGTATTATGT
601 AGTCTGTTTT TTATGCAAAA TCTAATTTAA TATATTGATA TTTATATCAT
651 TTTACGTTTC TCGTTCAGCT TT
```

#### 2 Decay of the autocorrelation of the gyration radius of the DNA minicircle

To sample the equilibrium behavior of the DNA minicircle, the trajectory length per MD replicate has been established upon the average decay of the autocorrelation of the gyration radius of the system:

$$R_g^2(t) = \frac{1}{N}\sum_{i=1}^{N}(\mathbf{r}_i(t) - \bar{\mathbf{r}}(t))^2 \quad (1)$$

with N the number of nucleotides and $\bar{\mathbf{r}}(t)$ the coordinates of the center of mass of the minicircle at time $t$: The characteristic decorrelation time of $R_g^2(\tau_d)$ is estimated as the first zero crossing of the autocorrelation function. Figure S1 depicts the autocorrelation of the gyration radius of the DNA minicircle for each superhelical density scenario, averaged over the independent replicates (shown in Figure S2).

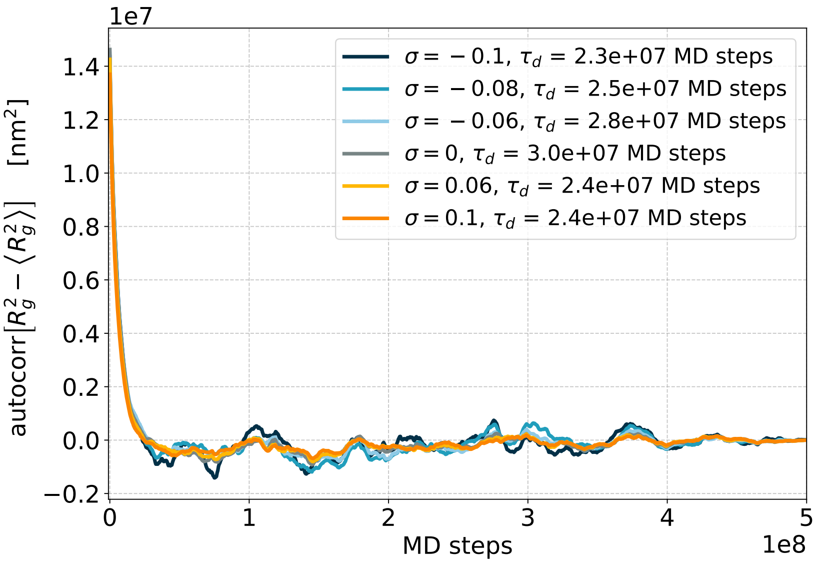


**Figure S1:** Autocorrelation of the gyration radius of the 672-bps DNA minicircle for each superhelical density scenario, averaged over ten independent replicates. Estimates of the characteristic decorrelation time of $R_g^2$ $(\tau_d)$, as the first zero crossing of the average autocorrelation function, are reported in the legend.

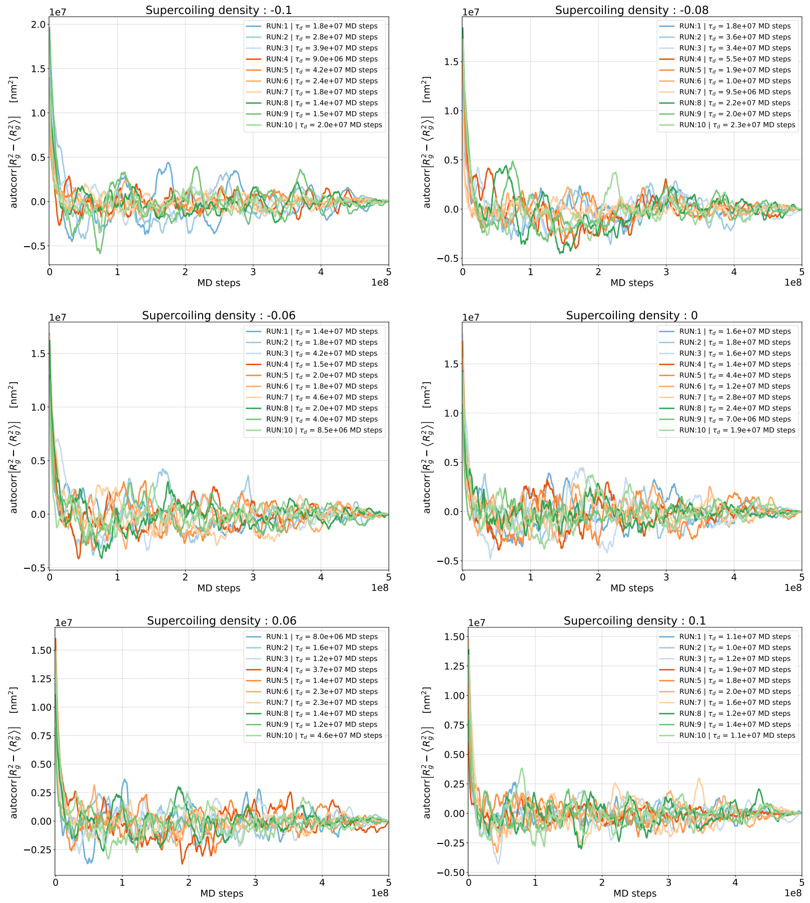


**Figure S2:** Autocorrelation of the gyration radius of the 672-bps DNA minicircle, for all independent MD replicates of each superhelical density scenario. Estimates of the decorrelation time of $R_g^2$, as the first zero crossing of the autocorrelation function, are reported in the legend.

### 3 Time evolution of the local writhe

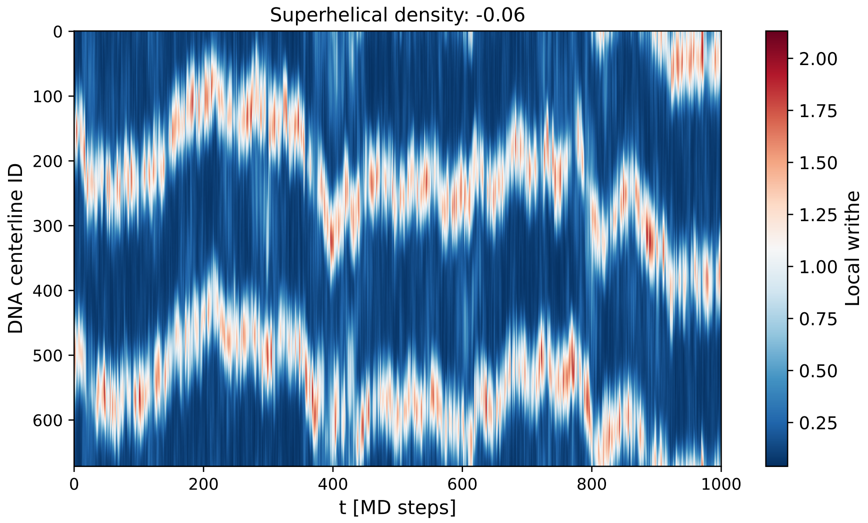


**Figure S3:** (Example) Time evolution of the local writhe along the centerline of a DNA minicircle at $\sigma = -0.06$, highlighting the dynamical behavior of the DNA structure and the topological motifs thereof.

### 4 Persistence and distribution of DNA denaturation bubbles

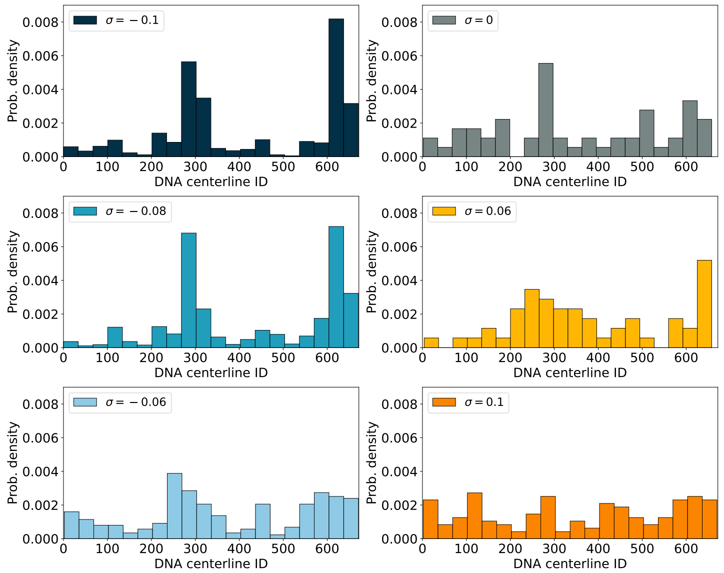


**Figure S4:** Probability density distribution associated with the likelihood for a nucleotide to host a DNA denaturation bubble, over all MD replicates of the 672-bps DNA minicircle at varying degrees of superhelical density. For the purpose of clarity, denaturation bubbles of size $B_s = 1$ have been excluded - in fact, they are distributed homogeneously along the DNA centerline.

| $\sigma$ | MD replicate (ID) | | | | | | | | | |
|---|---|---|---|---|---|---|---|---|---|---|
| | 1 | 2 | 3 | 4 | 5 | 6 | 7 | 8 | 9 | 10 |
| −0.1 | 88% | 100% | 96% | 100% | 96% | 94% | 89% | 90% | 98% | 100% |
| −0.08 | 42% | 48% | 57% | 52% | 60% | 38% | 47% | 51% | 45% | 53% |
| −0.06 | 26% | 27% | 28% | 29% | 27% | 29% | 29% | 27% | 26% | 29% |
| 0 | 19% | 17% | 20% | 19% | 18% | 18% | 18% | 19% | 20% | 17% |
| 0.06 | 17% | 16% | 20% | 19% | 15% | 16% | 16% | 20% | 18% | 19% |
| 0.1 | 18% | 21% | 21% | 19% | 16% | 17% | 18% | 16% | 19% | 19% |

**Table S1:** The percentage of frames over all independent MD replicates of the 672-bps DNA minicircle showing (at least) one denaturation bubble, at diverse degrees of superhelical density $\sigma$. A bubble is scored as the failure of hydrogen bonding interactions between base pairings belonging to the native conformation.

### 5 Time evolution of DNA denaturation bubbles

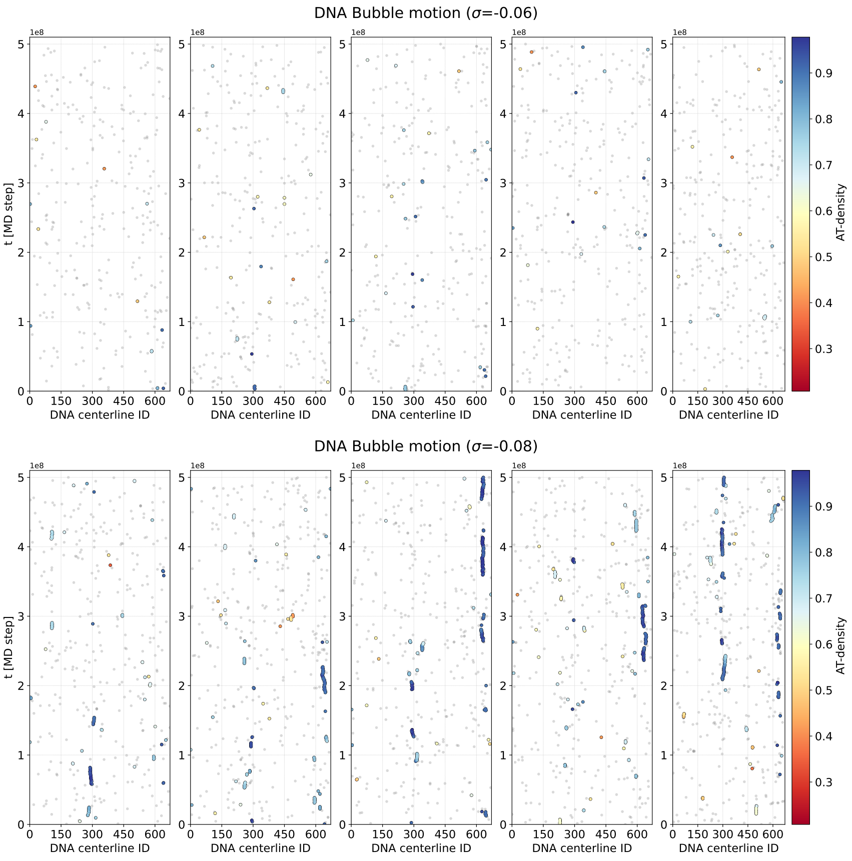


**Figure S5:** Time evolution of denaturation bubbles along the centerline of a DNA minicircle: Each data-point is defined by the center of mass of the bubble lying approximately about the $i$-th nucleotide. The color scale is associated with the local AT density (over a rolling average of 10 nucleotides) but for the denaturation events involving the failure of a single hydrogen bond, which are represented in gray for the purpose of clarity - as these defects are ubiquitously distributed along the DNA centerline. Here, the results of five independent MD simulations at superhelical density $\sigma = -0.06$ (**top**) and $\sigma = -0.08$ (**bottom**) are shown.

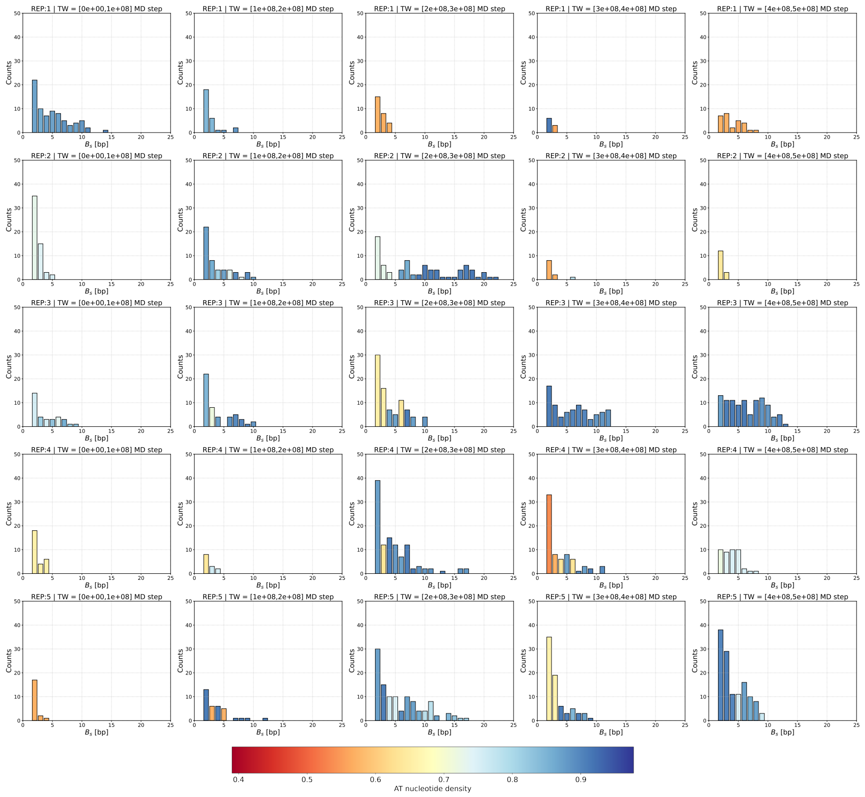


**Figure S6:** Histograms showing the distribution of DNA bubble sizes within time windows of $1 \times 10^8$ MD steps, along a selection of MD replicates in the $\sigma = -0.08$ scenario (corresponding to those shown in the upper panel of Figure S5. A color code has been defined upon the statistical mode of the distribution of AT-density values co-localizing with a specific $B_s$ shape. For the purpose of clarity, all events involving the failure of single hydrogen bonding interactions have been neglected.

#### 6 Structural characterization of synthetic bubbles

| $B_s$ [bps] | Residues involved with the bubble [idx] | $B_{CM}$ [idx] |
|---|---|---|
| 1 | 92 | 92 |
| 2 | 91-92 | 92 |
| 5 | 88-92 | 90 |
| 10 | 83-92 | 88 |

**Table S2:** The setups (shape, location) employed in the characterization of synthetic DNA bubbles. $B_s$ and $B_{CM}$ define the size and center of mass of the bubble respectively.

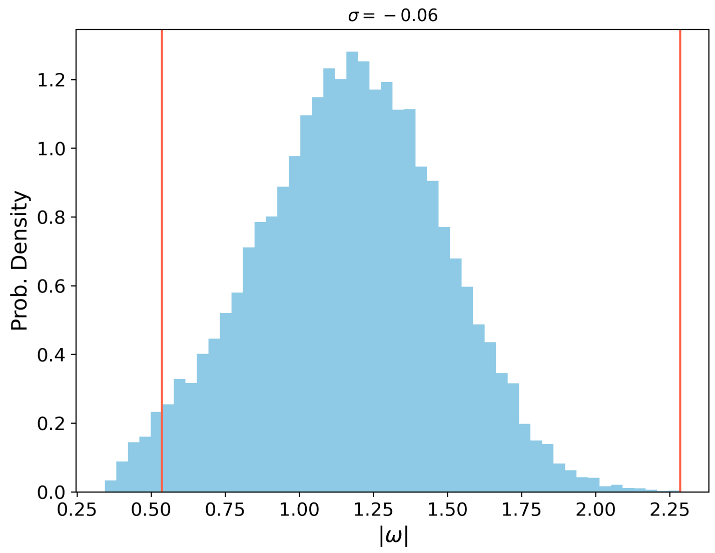


**Figure S7:** Distribution of the local writhe values associated with the apical loop regions of a 672-bp DNA minicircle at equilibrium and superhelical density $\sigma = -0.06$. The vertical red lines identify the lower and upper bounds of the gray area shown in Figure 9 of the main text - namely, $\langle\omega\rangle - 2\text{Var}[\omega]$ and $\max(\omega)$.
